# Prohibitin protects against cigarette smoke extract-induced cell apoptosis in cultured human pulmonary microvascular endothelial cells

**DOI:** 10.1101/2020.04.30.071589

**Authors:** Yating Peng, Zijing Zhou, Aiyuan Zhou, JiaXi Duan, Hong Peng, Ruoyun Ouyang, Yan Chen, Ping Chen

**Affiliations:** Department of Pulmonary and Critical Care Medicine, Changsha, Hunan, China, 410011; Institute of Respiratory Disease, Central South University, Changsha, Hunan, China, 410011; Hunan Centre for Diagnosis and Treatment of Respiratory Disease, Changsha, Hunan, China, 410011

**Keywords:** prohibitin, HPMECs, CSE, apoptosis, mitochondrial

## Abstract

Prohibitin is an evolutionarily conserved and ubiquitously expressed protein in eukaryocyte. It mediate many important roles in cell survival, apoptosis, autophagy and senescence. In the present study, we aimed to explore the role of prohibitin in cigarette smoke extract (CSE)-induced apoptosis of human pulmonary microvascular endothelial cells (HPMECs). For this purpose, HPMECs were trasfected with prohibitin and challenged with CSE. Our results showed that CSE exposure inhibited prohibitin expression in a dose-dependent manner in HPMECs. Overexpression of prohibitin could protect cell from CSE-induced injury by inhibiting CSE-induced cell apoptosis, inhibiting reactive oxygen species (ROS) production, increase mitochondrial membrane potential, increase the content of mitochondrial transcription factor A (mtTFA), IKKα/β phosphorylation and IκB-α degradation. CSE decreases prohibitin expression in endothelial cells and restoration of prohibitin expression in these cells can protect against the deleterious effects of CSE on mitochondrial and cells. We identified prohibitin is a novel regulator of endothelial cell apoptosis and survival in the context of cigarette smoke exposure.

## 1.0 Introduction

Active and passive smoking constitute a significantly public health problem. Cigarette smoke exposure is the leading cause of emphysema and triggers apoptosis within the lung parenchyma, resulting loss of the capillary and diminished gas exchange area. Lung parenchymal cells, especially endothelial cell apoptosis represents an early and a critical pathological process in the development of emphysema[1,2]. Studies conducted in our laboratory have revealed that CSE induce apoptosis in microvescular endothelial cells from lung[3,4]. CSE stimulation may generate ROS, induce DNA damage, leading to oxidative stress and subsequent endothelial cell apoptosis[5].

Prohibitin is a highly conserved and pleiotropic protein that is ubiquitously expressed in various compartments of eukaryocytes including mitochondrial, nucleus and plasma membrane. The main function of prohibitin in primary mammalian cells is to regulate nuclear transcription and mitochondrial integrity. Multiple prohibitin1(prohibitin) and prohibitin2 subunits forming ring-like prohibitin complexes in the inner mitochondrial membrane (IMM) and maintain the mitochondrial cristae structure. So, it is implicated in various cellular functions including cell apoptosis, autophagy, senescence, tumor suppression, transcription[6]. Increasing evidence revealed that ectopic expression of prohibitin protects cells and disease mouse models from oxidative stress injury through mitochondial mediated pathway. Moreover, prohibitin was reported to be abnormally downregulated in inflammatory bowel disease (IBD) and impeded tumor necrosis factor alpha (TNF-α)-induced epithelial barrier dysfunction and NF-κB activation [7].

While many studies have explored the functions of prohibitin, little is known about the function of this protein in the apoptosis of HPMECs induced by CSE. In COPD and non-COPD smokers, prohibitin protein and mRNA expression was significantly decreased compared to non-smokers and prohibitin expression levels are associated with the degree of airway obstruction[8]. In endothelial cells, the loss of prohibitin results in a block in electron transport at complex I, increased mitochondrial ROS generation, and cellular senescence that culminates in the loss of several endothelial functions including cell migration, cell proliferation, and blood vessel formation[9]. In the present study we aimed to validate the function of prohibitin, and illuminate the mechanisms of prohibitin in CSE-treated HPMECs apoptosis.

## 2.0 Materials and Methods

### 2.1 Cell culture

HPMECs were ordered from Science Cell company (Catalog Number: 3000) and were cultured in endothelial culture medium containing 10% FBS in a humidified incubator under 95% air, 5% CO_2_ at 37°C. HPMECs at 60% confluence were exposed to medium or CSE.

### 2.2 Preparation of CSE

CSE was prepared as previously reported with modification[10]. Briefly, one cigarette without filter was burned for 30sec and the smoke passed through 10 mL of endothelial cell medium using a vacuum pump. This 100% CSE was adjusted to a pH of 7.4 and filtered through a 0.22 μm filter (Millipore), and the CSE was diluted to the concentration of 0%, 1%, 1.5%, 2% and 2.5% in each and added to endothelial cells within 30 minutes of preparation. Cigarettes of a domestic brand were obtained from Changde Tobacco (Changde, Hunan, China).

### 2.3 Overexpression of prohibitin plasmid vectors

Briefly, the rat PHB1 cDNA was subcloned into the multiple cloning site of the shuttle plasmid pAdTrack-CMV. The purified recombinant plasmids were linearized and co-electroporated with pAdEasy-1 adenoviral backbone vector into Escherichia coli BJ5183. The complete adenovectors (Ad-PHB1) and empty adenovectors (Ad) were packaged by transfecting 293 cells, where viral particles were further amplified, purified, and titered.

### 2.4 Western blot assay

HPMECs were harvested and lysed with lysis buffer for 30 min on ice. After centrifugation for 30 min at 4°C, the supernatants were collected and protein concentration was determined by a bicinchoninic acid (BCA) protein assay kit (Pierce, USA). Equal amounts of protein were separated by sodiumdodecyl sulfate-polyacrylamide gel electrophoresis, electrotransferred to polyvinylidene difluoride (PVDF) membranes. Membrane was blocked with 5% BSA and incubated with the primary antibodies, followed by HRP-conjugated goat anti-rabbit secondary antibodies. Prohibitin and mtTFA, Phospho-IKKα/β (Ser176/180), IκB-α, NF-κB p65, IκB-α (Cell Signaling Technology), histone H3 (bioworld) and GAPDH (bioworld) antibody was used in this study.

### 2.5 Detection of Apoptosis

An Annexin V FITC Apoptosis Detection Kit II (BD Biosciences, San Jose, CA) was used to measure cell apoptosis. Briefly, HPMECs were collected and incubated with 5 μl Annexin V and 5 μl propidium iodide for 15 min in the dark at 25°C. The number of apoptotic cells was determined by flow cytometry. The early apoptotic cells were represented in the lower right quadrant of the fluorescence-activated cell sorting histogram. And late apoptotic cells and necrotic cells were represented in the upper right quadrant.

### 2.6 Caspase-3 activity assay

Caspase-3 activity was measured using caspase-3 activity assay kit (Beyotime, China) following the manufacturer’s protocol. Briefly, cells were resuspended in lysis buffer and centrifuged. Ac-DEVD-pNA and reaction buffer was added into cell lysis. After incubation at 37°C for 2h, absorbance was read on microplate reader at 405nm.

### 2.7 Detection of Mitochondrial membrane potential (MMP)

The HPMECs were washed with cold PBS and centrifuged at 1500rpm for 5min. 5×10^5^ cells were suspended in 0.5 ml cell culture medium and dyed by 0.5ml JC-1 dyeing fluid. Cells were incubated at 37°C for 20 minutes. After the incubation, cells were centrifuged at 1200 rpm for 3 min, and supernatants were discarded. Cells were washed by JC-1 dyeing buffer 2 times, suspended by 0.5 ml JC-1 dyeing buffer and analyzed by flow cytometry.

### 2.8 ATP contents assay

Cellular ATP contents were assessed by ATP Assay kit (S0026; Beyotime) according to the manufacturer’s instructions. Briefly, cells were rinsed and lysed using ATP lysis buffer on ice. Samples were collected and centrifuged at 12,000 rpm for 10 min at 4°C to acquire supernatant for further determination. Samples and ATP detection working dilution were added and luminescence activity was measured immediately using luminometer. Standard curve of ATP measure was made in each assay. Subsequently, the intracellular ATP contents were normalized by the protein contents in each sample.

### 2.9 ROS assay

The activity of ROS within the cell was determined using a standard 2’,7’-dichlorofluorescin diacetate (DCFDA) assay (Beyotime, China). Cells were incubated with DCFDA dye at 37°C for 20 min. Then, cells were observed under fluorescence microscopy and measured with fluorescence intensity.

### 3.0 Immunocytochemistry staining

After Leydig cells were treated in accordance with the above-described experimental design, Leydig cells on 6-chamber slides were fixed with 4% paraformaldehyde at 4°C for 15 min and then permeabilised with 0.2% Triton X-100 in PBS at room temperature for 15 min. e cells were then incubated with primary antibodies against mouse 8-hydroxy-2’-deoxyguanosine (8-OHdG) (1:200)(ab48508, 1:200 dilution; Abcam, Cambridge, United Kingdom) at 4°C overnight. Subsequently, 100 μL/well working solution of goat anti-mouse secondary antibodies (1:300) was added and incubated at room temperature for 90 min. After the cells were washed with PBS, they were stained with avidin-biotin-peroxidase complex visualised with DAB. The stained slides were photographed using an inverted microscope (Olympus, Tokyo, Japan) at 200 × magnification.

## 3. Statistical analysis

Data are reported as the mean ± SEM. Results were compared by 2-tailed Student’s t test for 2 groups and one-way ANOVA followed by the Tukey’s t test (2-tailed) for multiple groups. SPSS v16.0 (SPSS Inc., Chicago, IL) was used for analysis. Differences were considered statistically significant at P < 0.05.

## 4. Results

### 4.1. Effect of CSE on prohibitin expression

Western blot analysis of total cell extracts showed a steady decrease in prohibitin protein levels corresponding to 29kDa molecular mass when HPMECs were exposed to 0%, 1%, 1.5%, 2% and 2.5% CSE for 12 hours (Fig. 1A, B). The RT PCR analysis revealed that the mRNA levels of prohibitin significantly decreased in HPMECs as the concentration of CSE increased (Fig. 1C). We speculated that CSE induced cell apoptosis by down-regulating prohibitin.

**Figure 1.**
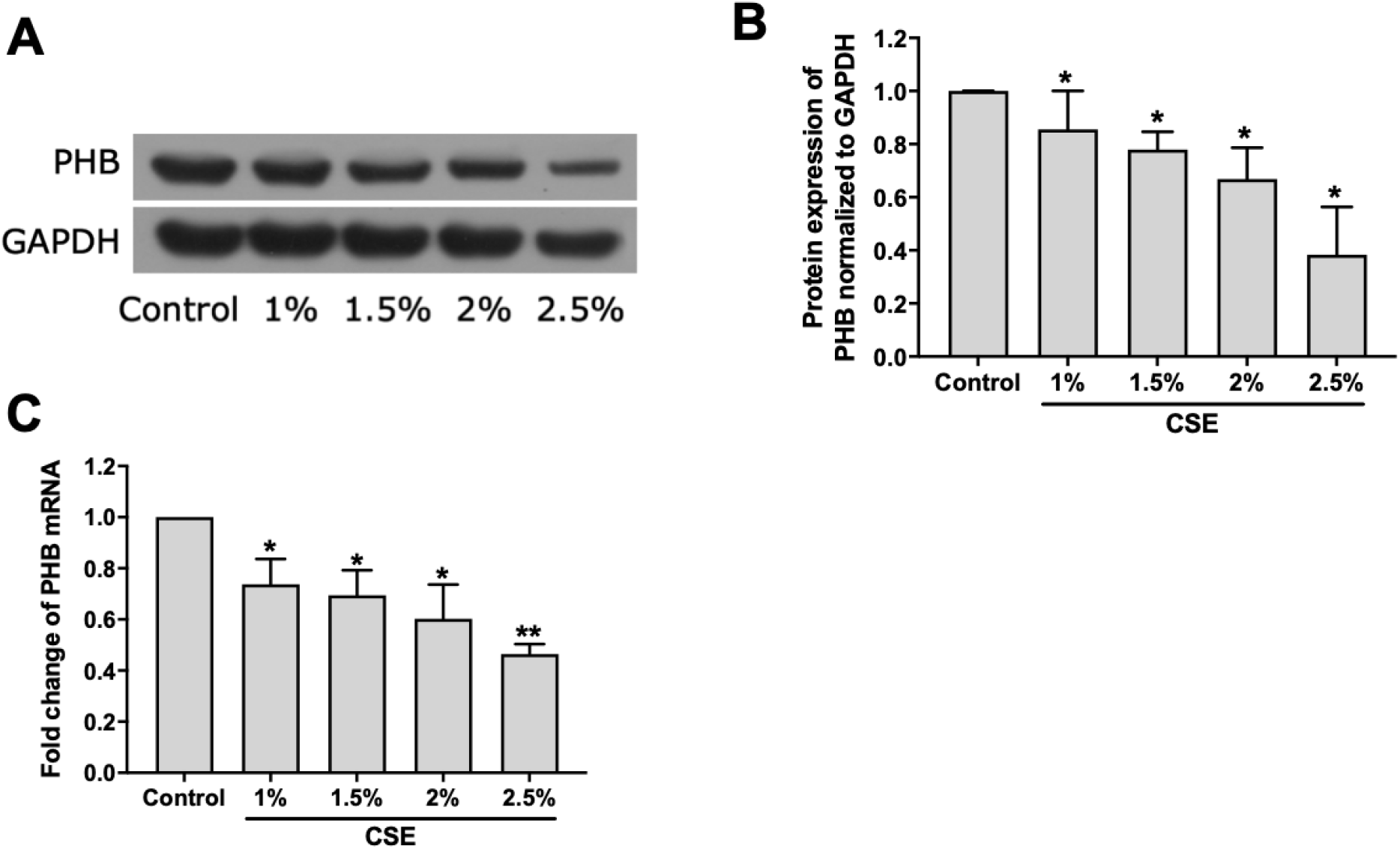
CSE induced dose-dependent downregulation of prohibitin in HPMECs. (A) Prohibitin protein levels were determined in HPMECs exposed to control medium or 1%, 1.5%, 2% and 2.5% CSE for 12 h. (B) Bar graphs represent the results from three independent experiments. (C) Relative mRNA expression were determined in HPMECs exposed to control medium or 1%, 1.5%, 2% and 2.5% CSE for 12 h. * P < 0.05 vs. control. ** P < 0.01 vs. control.

### 4.2. Effect of prohibitin overexpression on CSE mediated loss of MMP (Δψm) and ATP

Next, we explored the biological relevance of prohibitin upregulation in HPMECs after exposure to CSE. Protein expression of prohibitin remarkably elevated at 24 and 48 hours after transfection with prohibitin, while transfection with empty vector did not show any change in levels of prohibitin (Fig. 2A). MMP is a critical factor in maintaining the physiological function of mitochondria and is a major determinant of cell fate(Logan A, 2016). CSE exposure led to a significant perturbance of mitochondrial membrane potential compared with non-exposed HPMECs. Notably, prohibitin overexpression significantly attenuated CSE-induced loss of MMP (Fig. 2B, C). Energy production is a critical mitochondrial function. Treatment with CSE also significantly decreased ATP levels in HPMECs which were prevented by prohibitin overexpression (Fig. 2D). Our results indicate that prohibitin overexpression protectes cells from CSE promoting mitochondrial homeostasis.

**Figure 2.**
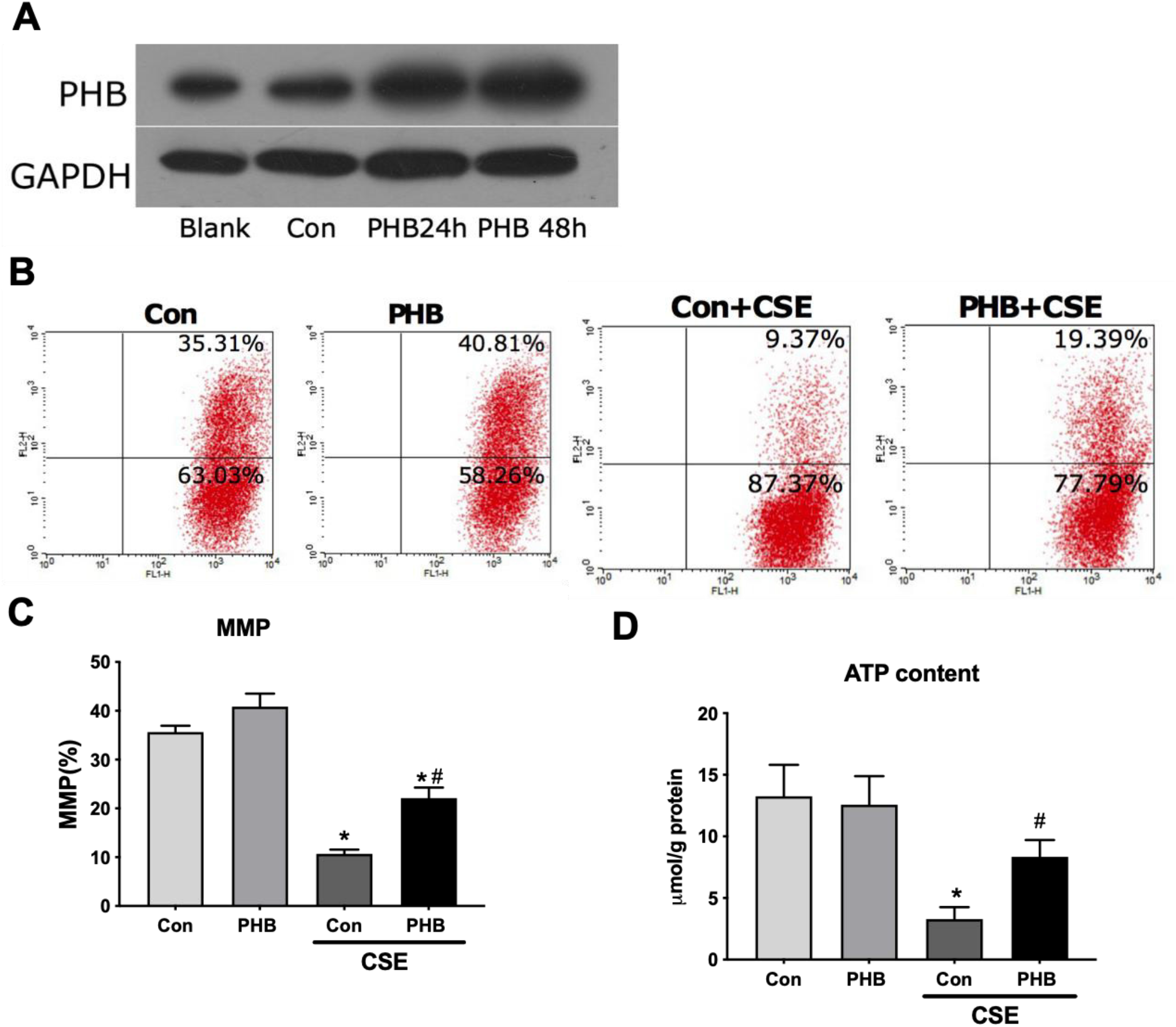
Prohibitin modulates changes in the Δψm and ATP content in HPMECs exposed to CSE. (A) HPMECs were transfected with the adenoviral PHB constructs for 2 h and cultured for 24 h or 48h. Then, PHB content in HPMECs was detected by Western blot method. (B) HPMECs were transfected with the adenoviral PHB constructs at a MOI of 2.3×10^9^ PFU/ml or empty vector for 2 h and cultured for 24 h. Then, HPMECs were cultured in fetal bovine serum-free media for 8 h and treated with 2.5% CSE for another 12 h. (B) Representative cytometry plots of cells incubated with JC-1 probe. (C) Bar graph demonstrates the levels of MMP at different groups. (D) ATP content analysis in HPMECs with indicated treatments. Bar graphs represent the results from three independent experiments. * P < 0.05 vs. Empty vector-transfected cells with control medium; # P < 0.05 vs. Empty vector-transfected cells with CSE.

### 4.3. Effect of prohibitin overexpression on CSE mediated ROS

It is well known that mitochondria are the primary source of cellular ROS, and ROS serve an important role in activation of apoptotic signaling. To determine whether there was a functional consequence of the loss of MMP and ATP, we examined intracellular ROS levels. Previous study have showed that CSE induced ROS production in endothelial cells.(Kim B S,2013) We confirmed that CSE exposure increased intracellular ROS levels and we further revealed that prohibitin overexpression inhibited intracellular ROS production under CSE treatment (Fig. 3A and B).

**Figure 3.**
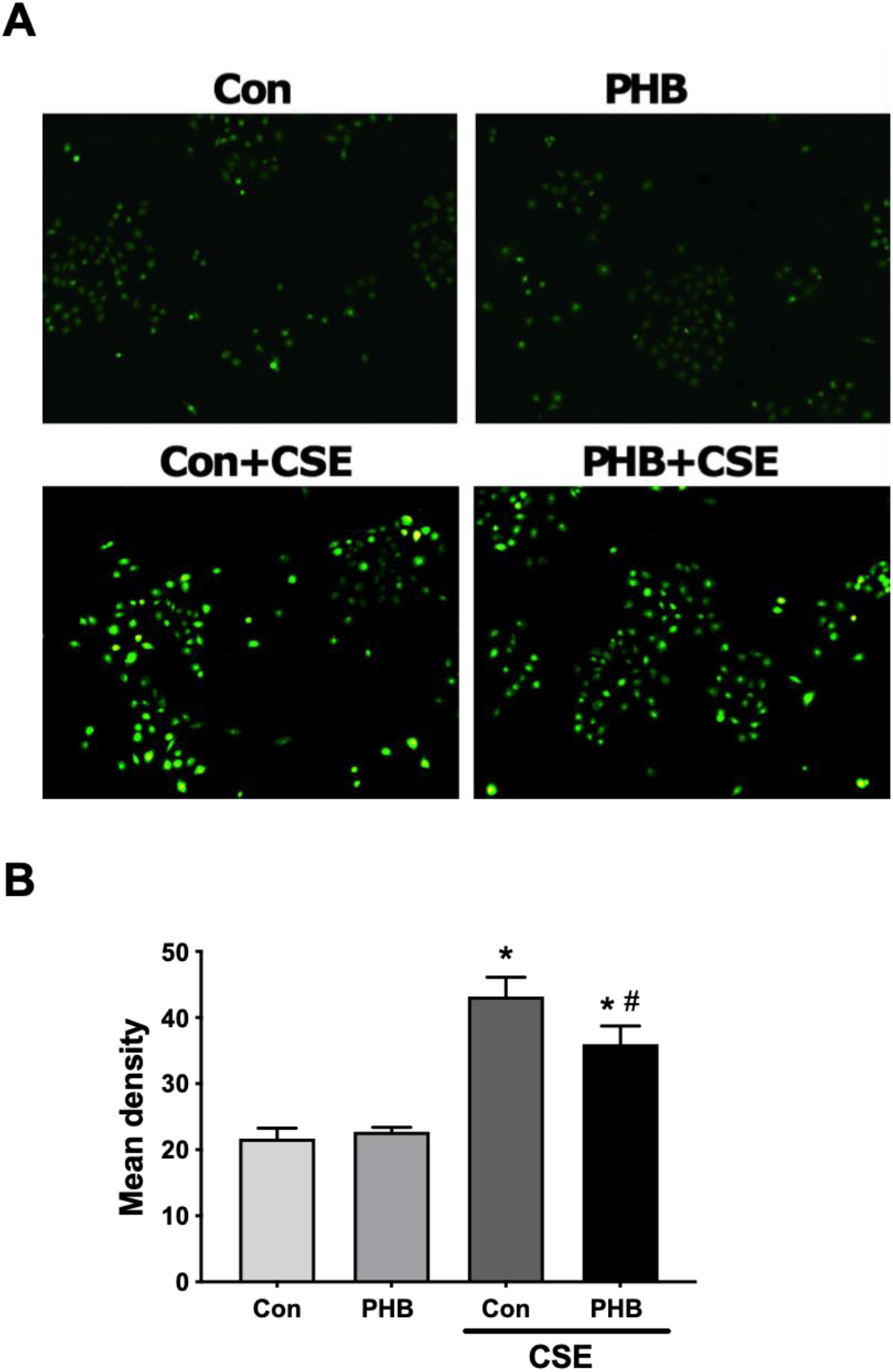
Prohibitin suppresses CSE-induced intracellular ROS in HPMECs. (A) Fluorescent photomicrographs of cellular ROS level in HPMECs transfected with prohibitin-expressing plasmid or vector and exposed to control medium or CSE. Original magnification×200. (B) Bar graph demonstrates the levels of ROS at different groups. * P < 0.05 vs. Empty vector-transfected cells with control medium; # P < 0.05 vs. Empty vector-transfected cells with CSE.

### 4.4. Effect of prohibitin overexpression on CSE mediated oxidant-induced DNA damage

Immunohistochemistry was done for the assessment of CSE induced oxidative DNA damage. In terms of 8-OHdG expression, CSE exposure group showed a remarkable amount of 8-OHdG both in the nuclear and cytoplasm. The results indicated that CSE could seriously damage DNA and prohibitin overexpression mitigated the oxidative DNA damage associated with it (Fig. 4A and B).

**Figure 4.**
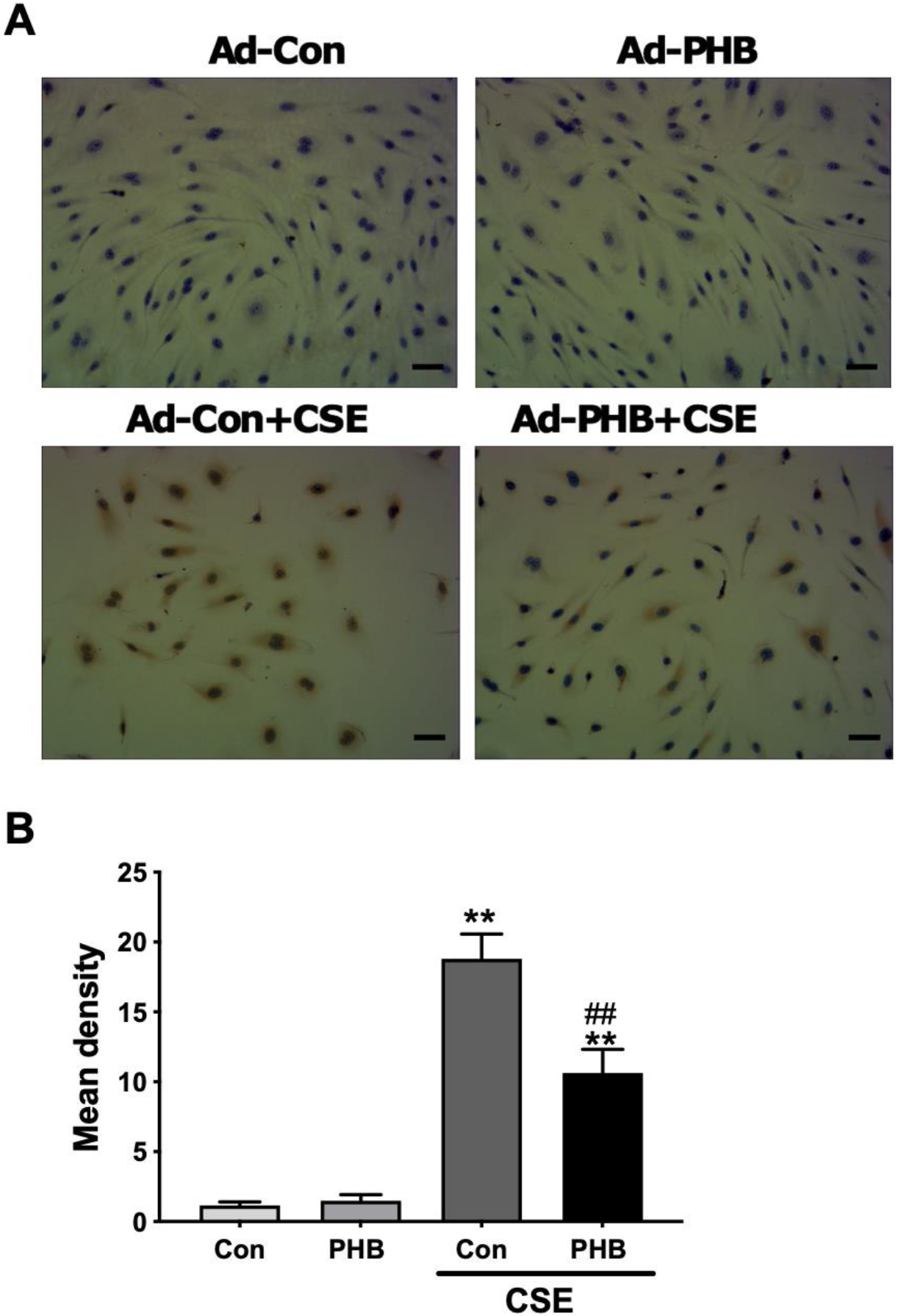
Prohibitin suppresses CSE-induced oxidative damage in HPMECs. (A) Immunohistochemistry photomicrographs of 8-OHdG expression in HPMECs transfected with empty vector or prohibitin-expressing plasmid and exposed to control medium or CSE. Original magnification×400. (B) Bar graph demonstrates the levels of 8-OHdG at different groups. * P < 0.05 vs. Empty vector-transfected cells with control medium; # P < vs. Empty vector-transfected cells with CSE.

### 4.5. Effect of prohibitin overexpression on CSE induced apoptosis

As demonstrated by our previous experiments, CSE treatment significantly triggered apoptotic cascade response in HPMECs(Zong D, 2018). By using flow cytometric quantitation of annexin V-propidium iodide staining, we found overexpression of prohibitin prevented apoptosis of HPMECs induced by CSE (Fig. 5A,B). Next, we asked whether caspase 3-mediated cell death existed in CSE-treated HPMECs and this process was weakened by prohibitin overexpression. (Fig. 5C).

**Figure 5.**
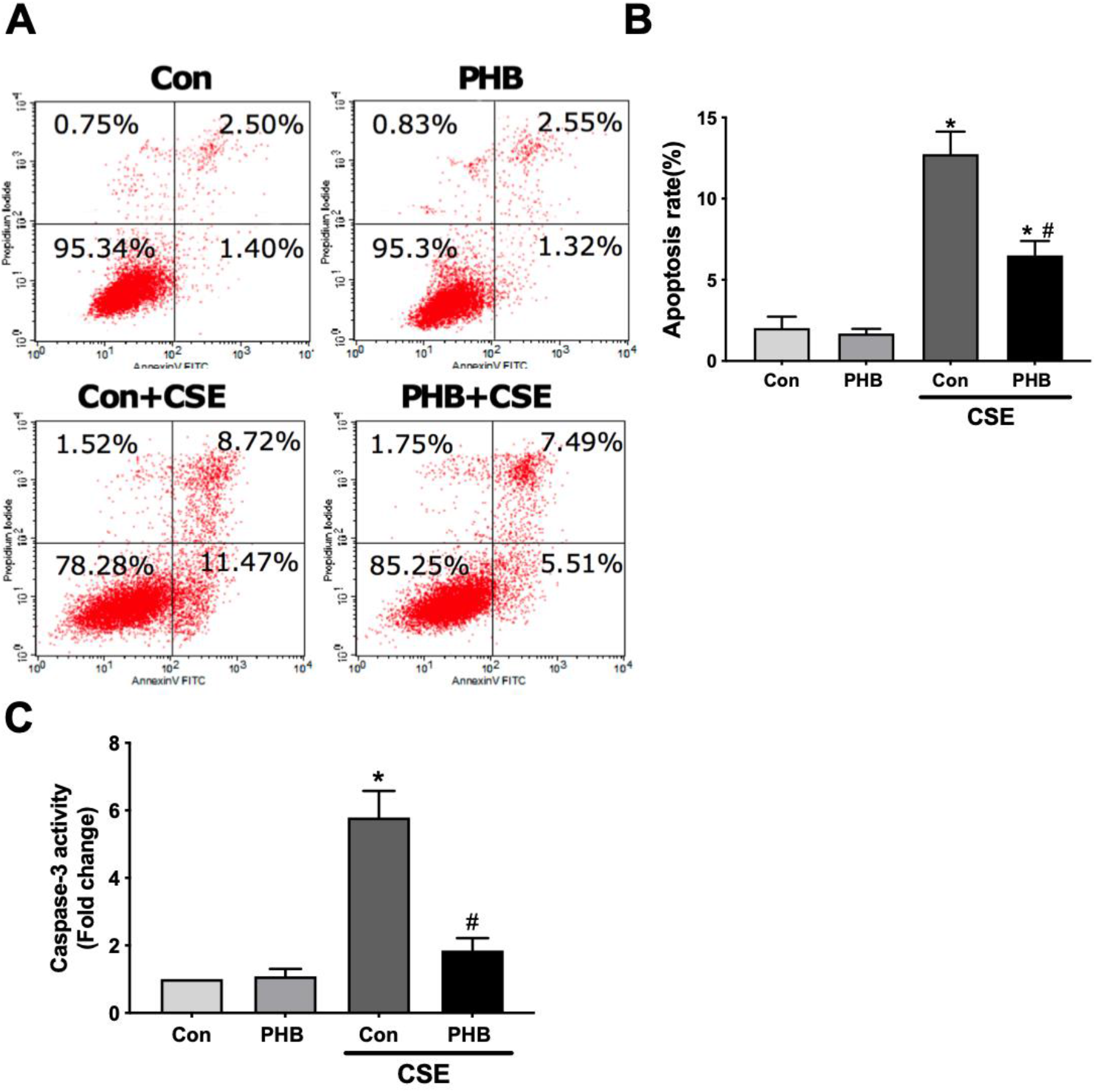
Prohibitin protects HPMECs from CSE-induced apoptosis. (A)The apoptosis rate of HPMECs were tested by flow cytometry. (B) Bar graph demonstrates the apoptosis rate at different groups. (C) Measurement of caspase-3 activity in HPMECs transfected with prohibitin-expressing plasmid or empty vector before exposure to control medium or CSE. The results are expressed as fold change relative to empty vector-transfected cells treated with control medium. ** P < 0.01 vs. Empty vector-transfected cells with control medium; ## P < 0.01 vs. Empty vector-transfected cells with CSE. Scale bar, 20 μm.

### 4.5. Effect of prohibitin overexpression and CSE on NF-κB activation

We found CSE induced decreased expression of mTFA protein and prohibitin overexpression increase mtTFA protein expression. CSE-induced IKKα/β protein phosphorylation, was attenuated by prohibitin overexpression. CSE-induced IκB-α degradation, indicated by the markedly decrease of IκB-α in the cytoplasmic fraction of HPMECs, was also inhibited by prohibitin overexpression (Fig. 6A, B). Meanwhile, Nuclear and cytosolic extracts were assayed for p65 expression. However, we did not detect any significant change in the expression of nuclear and cytosolic NF-κB p65 expression. Altogether, these results suggest that Prohibitin inhibits CSE-induced mitochondrial fission by modulating the phosphorylation of IKKα/β and IκB-α degradation (Fig. 6 C, D).

**Figure 6.**
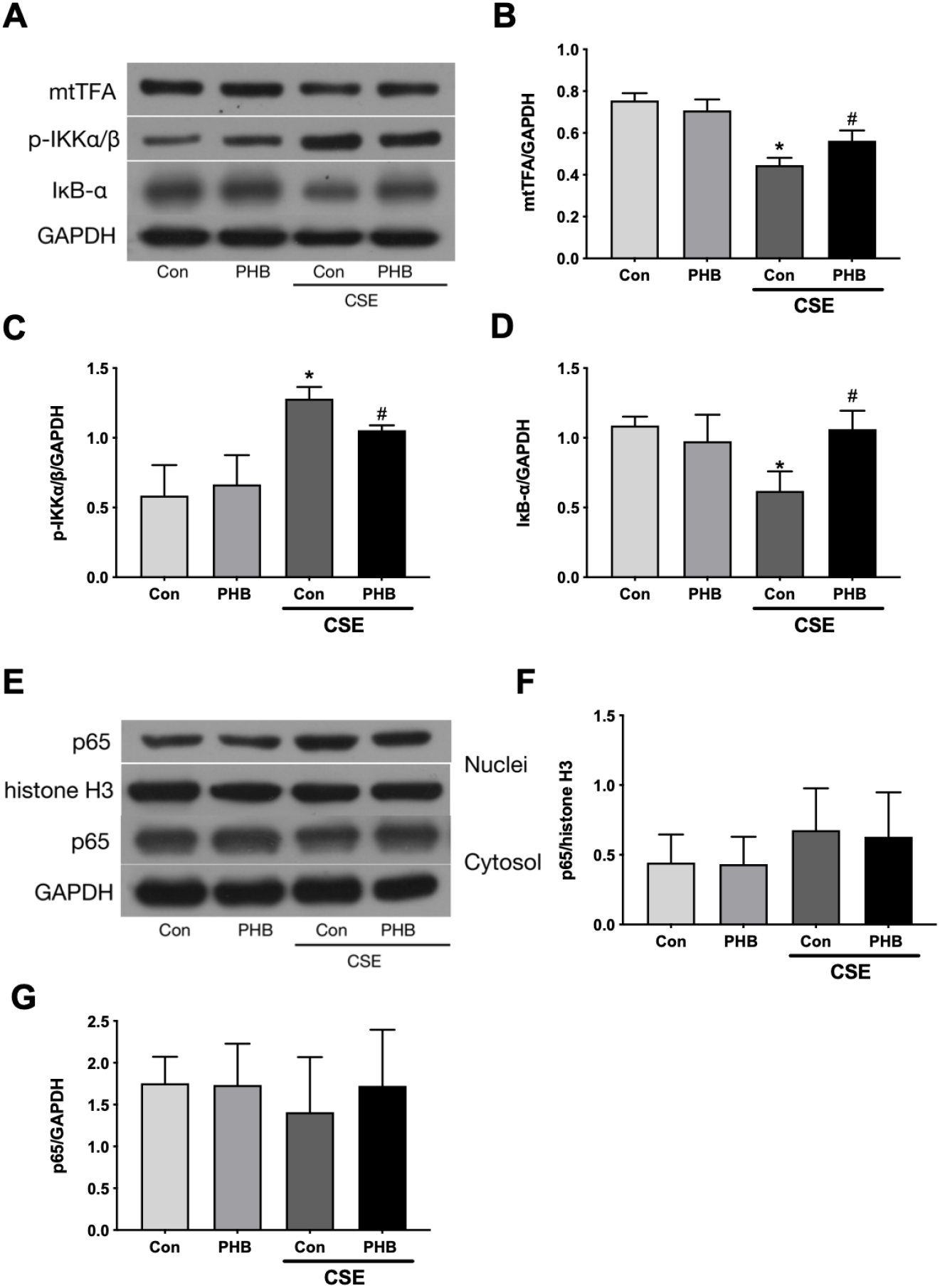
Prohibitin changed protein expression in HPMECs under CSE. (A) Western blot analysis of mtTFA, Phospho-IKKα/β, and IκB-α protein in HPMECs with indicated treatments. (B, C, D) Quantitative intensity data of the mtTFA, Phospho-IKKα/β, and IκB-α protein bands normalized to GAPDH. (E) Western blot analysis of Nuclear and cytosolic p65 expression in HPMECs with indicated treatments. (F, G) Quantitative intensity data of the p65 protein bands normalized to histone H3 or GAPDH. * P < 0.05 vs. Empty vector-transfected cells with control medium; # P < 0.05 vs. Empty vector-transfected cells with CSE.

## 5. Discussion

In the present study, we have investigated the protective ability of prohibitin overexpression in CSE exposure associated mitochondrial dysfunction, ROS generation, oxidative DNA damage and apoptosis in HPMECs.

CSE may induce apoptosis in bronchial epithelial cells, endothelial cells and human airway smooth muscle cells (ASMCs)[11–14] CSE may cause injury in mitochondrial morphology and function and induce apoptosis via an intrinsic apoptotic pathway, which involves mitochondrial fragmentation and the disruption of mitochondrial membrane integrity, inhibit of mitochondrial respiration and release of cytochrome c, as well as changed expression levels of pro-apoptotic and anti-apoptotic molecules[13,15–18]. A previous study has demonstrated that prohibitin decrease the apoptosis of endothelial cells in response to glyLDL[19]. Another study has shown that high prohibitin tumor expression is associated with poorer overall survival in patients with non-small cell lung cancer (NSCLC) and systemic delivery of PHB1 siRNA drastic inhibit tumor growth[20]. These results suggest a pro-survival activity prohibitin.

Previous studies have shown a negative effect of oxidative stress like hydrogen peroxide (H_2_O_2_) on the expression of prohibitin in human renal proximal tubule epithelial cell line and adult retinal pigment epithelial cell line-19[21,22]In agreement with these studies, we discovered that prohibitin expression is significantly reduced in endothelial cells in a CSE dose-dependent manner. The mRNA levels of prohibitin significantly decreased at 2.5% CSE for 12 hr, and the protein levels of prohibitin also decreased to some extent at 2.5% CSE. Therefore, the cells exposed for 2.5% CSE and 12 h were used for subsequent experiments. Furthermore, our results showed that the mRNA levels of prohibitin were reduced in endothelial cells exposed to CSE, indicating a defect in the transcriptional regulation of prohibitin. However, as microRNA may directly targeted prohibitin and phosphorylation of prohibitin plays a pivital role in cell survival[23,24], a post-translational deimination of prohibitin in CSE-induced apoptosis in HPMECs also can’t be excluded.

We then investigated the contribution of prohibitin to mitochondrial function under CSE treatment. The term mitochondrial dysfunction is referred to as a complex display of cellular events. Mitochondrial injury causes change in mitochondrial morphology, impaired oxidative phosphorylation, reduced membrane potential, altered metabolic activity, increased mitochondrial superoxide levels including alterations in intracellular Ca^2^+ flux[25,26]. We observed a protective effect of prohibitin overexpression on the decline in ATP levels and MMP in HPMECs. Prohibitin is a key regulator of mitochondrial quality control[27]. As its uniquely complex structure in mitochondrial localization, prohibitin exert protection function by maintain mitochondrial structure and morphology. Importantly, prohibitin complex is essential for mitochondrial biogenesis and degradation, and responding to mitochondrial stress[27]. As demonstrated by other studies, prohibitin overexpression prevents mitochondrial fragmentation, preserved mitochondrial respiratory function, attenuated mitochondrial complex I oxidative degradation of cells exposed to oxidative stress[28]. Our data provide the evidence that CSE caused mitochondrial depolarization, decreased energy production and prohibitin modulation affects mitochondrial function in this type of the cell under CSE treatments.

In the current study, we also verified that CSE increased ROS production and oxidative DNA damage in HPMECs. The general idea of the interplay between depolarization of the mitochondrial membrane and ROS production is that, upon mitochondrial damage or dysfunction, mitochondrial pores open and allowing the influx of potassium and calcium cations, thereby depolarizing the mitochondrial membrane, which in turn induces ROS production and release. Meanwhile, excessive ROS directly causing the collapse of MMP and depletion of ATP, which later activates a series of signaling pathways that induce apoptosis[29]. Several studies showed prohibitin acts as a coactivator for ARE-dependent gene expression and promotes endogenous antioxidant defense components under oxidative stress[30–32]. Prohibitin overexpression significantly reduced the level of ROS within HPMECs under CSE, suggesting that reduction of oxidative stress is one major mechanism of prohibitin-mediated endothelial cell protection. The oxidative damage in DNA by cigarette smoke is either caused directly, or through the generation of ROS[33]. Prohibitin interact with the Nicotinamide adenine dinucleotide hydrogen (NADH) dehydrogenase protein complex, which is essential for oxidoreductase activity and DNA repair within cells[34,35]. Hence, prohibitin may act as an anti-oxidative agent by inhibiting CSE-induced oxidative damage and oxidative DNA damage.

mtTFA, a nucleus-encoded protein, regulates the transcription and replication of mtDNA and maintains mtDNA copy number[36]. Massive apoptosis was found in mtTFA knockout embryos and in the heart of the tissue-specific mtTFA knockouts[37], and the respiratory chain deficiency caused by mtTFA knockout predisposes cells to apoptosis[37]. TFAM overexpression inhibits mitochondrial ROS production and reduces oxidative stress to mtDNA[38,39]. As demonstrated by our previous studies, in COPD patients with squamous cell lung cancer, and the level of mtTFA protein in lung tissue negatively correlates with pulmonary vascular endothelial apoptotic index and smoking index[40]. Expression of mtTFA mRNA and protein was downregulated in CSE-treated HUVECs as a consequence of hypermethylation of the mtTFA promoter[40]. Prohibitin subunits have been involved in the organization and stability of mitochondrial nucleoids together with mtDNA-binding proteins, mtTFA, and mitochondrial single-stranded DNA binding protein (mtSSB). In HeLa cells, prohibitin regulates copy number of mtDNA by stabilizing mtTFA protein, probably in a chaperone-like fashion[41]. Our study also demonstrated that mtTFA is a downstream effector of prohibitin that modulates mitochondrial function in CSE-exposed HPMECs.

Since a previous study showed that NF-κB activation is considered a central event in endothelial cells’ response to inflammatory stimuli. We also found CSE exposure markedly increased IKKα/β phosphorylation and IκB-α degradation in HPMECs. Interstingly, prohibitin did not only attenuate CSE-induced phosphorylation of IKKα/β but also restore the level of IκB-α. However, the mechanism through which prohibitin affects NF-κB remains to be established. It has been shown that ROS can both activate and repress NF-κB signaling in a phase and context dependent manner[42]. While overexpression of mtTFA significantly inhibited rotenone-induced mitochondrial ROS generation, mitochondrial DNA Damage and the subsequent NF-κB nuclear translocation[43]. Interestingly, prohibitin overexpression decrease NF-κB transcription activity, even after the proinflammatory cytokine LPS or TNF-α stimulation [7,44].

There are several open questions worth future study. First, we didn’t explore the phosphorylation or other post transcriptional modification state of prohibitin, since these epigenetic changes is far more important than relative expression for cell function. Second, it is interesting to explore if there is protein–protein interaction between mtTFA and prohibitin. Third, if changed NF-κB and mtTFA signal could insult the protective effect of prohibitin overexpression on HPMECs under CSE is unclear.

Although the molecular pathways underlying the prohibitin-mediated protection on HPMECs under CSE treatment are not completely understood, our results indicated that prohibitin could serve as a mitochondrion-targeting agent. In conclusion, our data suggested that prohibitin protects the HPMECs from CSE-induced apoptosis, and mitochondrial apoptosis pathway was involved in this process.

## Data Availability

The data used to support the findings of this study are available from the corresponding author upon request.

## Acknowledgments

This project carried out with the support of the National Key Clinical Specialist Construction Projects ((2012)No. 650) and National Natural Science Foundation of China (81370143,81170036).

